# Calcium-permeable AMPAR in hippocampal parvalbumin-expressing interneurons protect against memory interference

**DOI:** 10.1101/2025.04.29.651201

**Authors:** Matthew Cooper, Mina Moniri, Mohammed Abuelem, David M Bannerman, Edward O Mann

**Author notes:** Corresponding authors &.

## Abstract

Parvalbumin-expressing (PV+) interneurons exert exquisite control over spike output, and plasticity in these inhibitory circuits may be important for maintaining network stability in learning and memory. PV+ interneuron recruitment is primarily mediated by GluA2-lacking Ca^2+^-permeable AMPA receptors (CP-AMPAR), which support anti-Hebbian plasticity. However, the functional significance of CP-AMPAR-mediated plasticity remains unknown. Using a viral approach to artificially express the GluA2 subunit in hippocampal PV+ interneurons, we replaced CP-AMPAR with GluA2-containing receptors, and in doing so reduced synaptically-evoked Ca^2+^ transients and anti-Hebbian plasticity. Transfection of hippocampal PV+ interneurons with GluA2 resulted in delay-dependent spatial working memory deficits which increased across trials per session, and impaired reversal learning in the Morris water maze but not initial acquisition. Our data suggest that loss of CP-AMPAR-mediated plasticity in these cells leads to proactive interference, revealing a significant role for dynamic recruitment of PV+ interneurons in the segregation of memories and accurate memory retrieval.

**Highlights:**

- Viral expression of GluA2 in PV+ interneurons alters synaptic AMPA receptor profile
- GluA2 overexpression in PV+ cells reduces synaptic Ca^2+^ transients and plasticity
- Upregulating GluA2 in PV+ cells causes delay-dependent deficits in working memory
- Acquisition of reference memories is preserved, but reversal learning is impaired

## Introduction

Plasticity in inhibitory circuits is believed to be important in preserving excitatory-inhibitory balance, maintaining network stability and the segregation of competing memory traces^1-3^. Fast-spiking parvalbumin-expressing (PV+) interneurons are critical for controlling spike timing and generating patterns of rhythmic activity in hippocampal circuits^4^. Recruitment of these cells is predominantly mediated by GluA2-lacking AMPA receptors, which are Ca^2+^-permeable and display polyamine-dependent inward rectification^5,6^. These properties of Ca^2+^-permeable AMPA receptors (CP-AMPAR) enable anti-Hebbian plasticity at excitatory synapses onto PV+ cells, whereby pre-synaptic inputs associated with post-synaptic membrane hyperpolarisation result in Ca^2+^ influx and synaptic potentiation^7^, and vice versa^8,9^. However, the contribution of anti-Hebbian plasticity in hippocampal PV+ interneurons to network function and behaviour remains to be established.

To reduce CP-AMPAR expression in specific cell types, a viral construct was generated for driving Cre-dependent expression of the mouse GluA2 subunit and transfected into the hippocampus of PV-Cre-expressing mice. This approach reduced functional CP-AMPAR expression in PV+ interneurons without abolishing synaptic drive, leading to impairment of anti-Hebbian plasticity and increased interference between competing hippocampal memories.

## Results

### Transfection of Cre-expressing PV+ cells drives expression of GluA2 construct

The disruption of Ca^2+^-permeability by GluA2 is due to a post-transcription RNA edit, resulting in a glutamine-to-arginine (Q/R) substitution in the channel pore^10^. Therefore, we utilised a construct in which cDNA encoding the Q/R-edited, flop splice variant of the GluA2 subunit, with an AU1 peptide tag in its N terminus, was flanked by loxP sites and positioned downstream of a synapsin (hSyn) promoter. The construct was packaged in an rAAV8 adeno-associated virus and injected into the hippocampus of mice double homozygous for PV+ cell-specific Cre recombinase expression (PV-Cre) and Cre-dependent tdTomato expression (Ai9). Immunofluorescence against the AU1 epitope revealed diffuse immunoreactivity throughout the neuropil of injected hippocampus (Figure 1A&B). At a cellular level, anti-AU1 immunoreactivity was localised to neurons co-expressing tdTomato (i.e. PV+ cells, Figure 1C), and was restricted to extranuclear cytoplasm and visible dendritic regions, consistent with the predominantly dendritic localisation of AMPAR^11^. Histology indicated successful uptake and expression of the construct by PV+ interneurons across the different hippocampal subfields.

**Figure 1.**
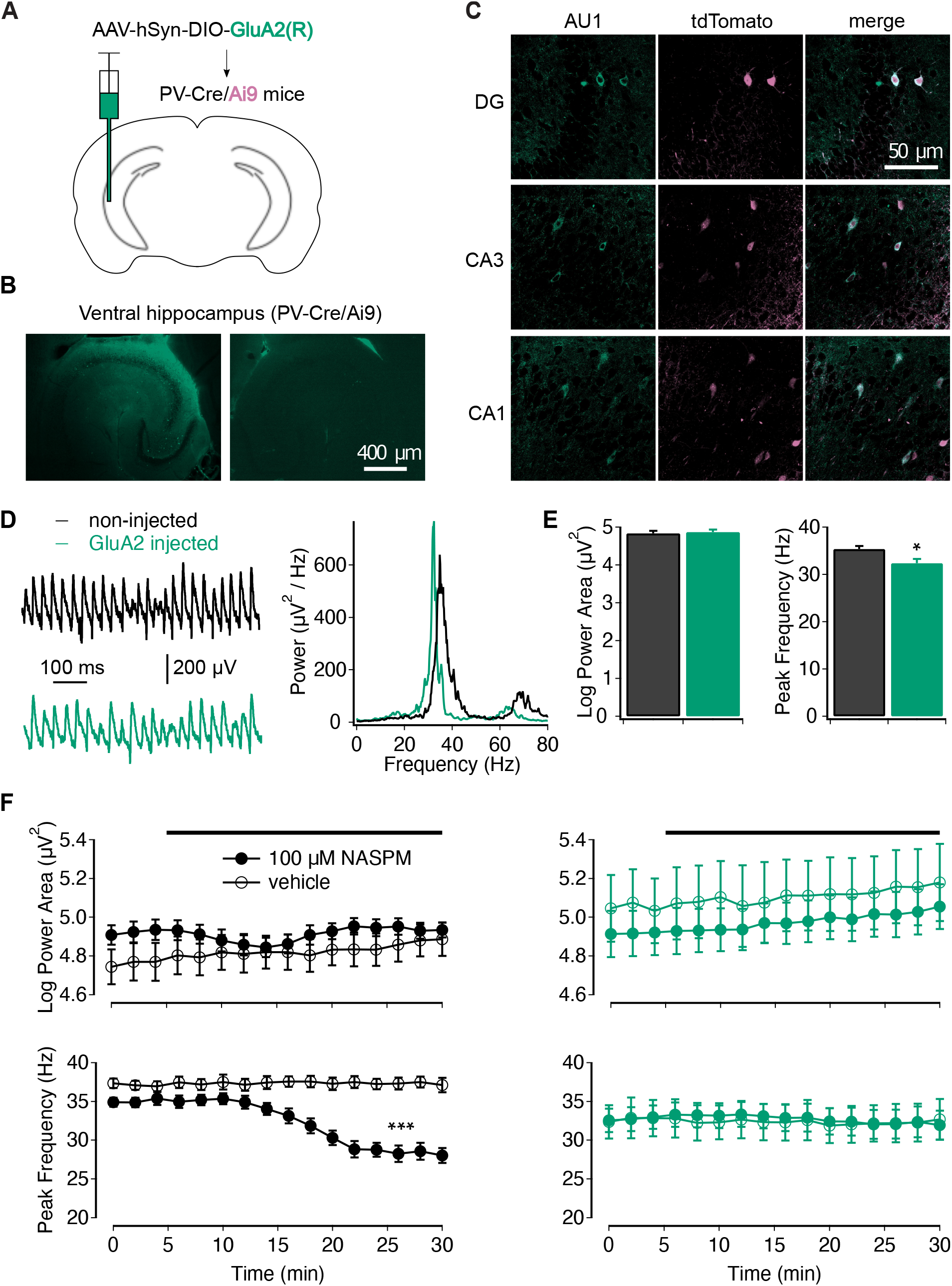
Overexpression of GluA2 in PV+ interneurons alters network pharmacology. (A) Schematic of unilateral viral injections into ventral hippocampus. (B) Anti-AU1 immunoreactivity in injected hemisphere (left) and uninjected hemisphere (right) in ventral hippocampus of PVCre/Ai9 mice. (C) Anti-AU1 immunofluorescence (left), tdTomato expression (middle) and merged images (right) in different subfields of ventral hippocampus. (D) Representative local field potential recordings (left) and power spectral density plots (right) of carbachol-induced gamma-frequency oscillations in ventral hippocampus of non-injected and GluA2-injected mice. (E) Expression of GluA2 in PV+ interneurons had no significant effect on the power of carbachol-induced oscillations (t = 0.35, p = 0.73, n = 33 for non-injected, n = 27 for GluA2-injected), but was associated with a significant reduction in peak frequency (t = 3.03, p = 0.004; unpaired t tests). (F) Effects of 100 µM NASPM on carbachol-induced oscillations in slices from non-injected (left) and GluA2-injected mice (right), measured in terms of power (top; F_(1,35)_ = 0.37, p = 0.55 for main effect of drug; F_(1,35)_ = 1.09, p = 0.30 for interaction; two-way ANOVA on change in power at 10 minutes post-treatment) and frequency (bottom; F_(1,35)_ = 40.00, p = 0.006 for main effect of drug; F_(1,35)_ = 55.84, p = 0.001 for interaction; two-way ANOVA on change in peak frequency at 10 minutes post-treatment). A reduction in peak frequency in slices from non-injected controls (t = 5.59, p < 0.001, n = 9 & 11 for vehicle and NASPM treatments, respectively; unpaired t test), was prevented by GluA2 transfection (t = 0.32, p = 0.75, n = 7 & 12 for vehicle and NASPM treatments, respectively; unpaired t test). *p < 0.05; ***p < 0.001. Values represent mean ± SEM.

### Viral expression of GluA2 in PV+ interneurons alters network pharmacology

Synaptic feedback loops between CA3 pyramidal neurons and PV+ interneurons support gamma-frequency (γ) oscillations^12,13^, and genetic knockout of AMPAR subunits in PV+ interneurons causes reductions in hippocampal γ power^14^. In order to explore the functional consequences of GluA2 expression in PV+ cells, we first examined cholinergically-induced γ oscillations in ventral hippocampal slices^15^, including baseline properties and sensitivity to CP-AMPAR-selective antagonists. Oscillations could be generated in slices from both non-injected control mice and GluA2-injected animals (Figure 1D). However, whilst γ power recorded from slices in the two groups did not differ significantly, oscillations from GluA2-injected mice were of significantly lower frequency than those from control slices (35.37 ± 0.6 Hz vs 32.35 ± 0.9 Hz; Figure 1E). Slices were subsequently exposed to the selective CP-AMPAR antagonist, 1-naphthylacetyl spermine (NASPM; 100 µM). In control slices, NASPM caused a transient reduction in γ power, followed by a sustained decrease in frequency (Figure 1F), possibly reflecting a switch to a different interneuronal circuit^16^. In contrast, these effects of NASPM were absent in slices from GluA2-injected animals (Figure 1F). These results reveal that overexpression of GluA2 in hippocampal PV+ interneurons does not affect the ability to generate carbachol-induced γ oscillations but eliminates the effects of a CP-AMPAR-selective antagonist on oscillations, demonstrating a functional effect of the viral manipulation on synaptic AMPAR.

### Expression of GluA2 reduces synaptically-evoked dendritic Ca^2+^ signals and anti-Hebbian plasticity

The inclusion of the GluA2 subunit should reduce inward rectification of synaptic AMPAR and impair AMPAR-mediated Ca^2+^ influx. A significant reduction in inward rectification was observed for evoked excitatory postsynaptic currents (EPSCs) in PV+ interneurons from GluA2-injected animals (U = 115, p = 0.01 c.f. non-injected controls; Mann-Whitney U test; Figure S1). There were no significant differences detected in spontaneous EPSC properties, although there was a correlation between the decay constant of spontaneous EPSCs and the rectification of evoked EPSCs (r = 0.73, p = 0.008; Figure S2), suggesting that increasing GluA2 expression leads to slower synaptic responses^17^. To explore further the functional consequences of changes in AMPAR expression, we imaged synaptically-evoked Ca^2+^ signals in CA1 PV+ cells in ventral hippocampal slices following intracellular loading of Oregon Green 488 BAPTA-1 (OGB-488; 300 µM). The ‘threshold’ stimulation amplitude was set as the minimum intensity necessary to reliably elicit a 50-100 pA EPSC, and responses were recorded between 100-300% of threshold (Figure 2A). Application of 100 µM NASPM significantly reduced the size of these Ca^2+^ signals (Figure 2A&B). The effects of NASPM were more pronounced at strong stimulation amplitudes, with significant effects observed at 200 and 300% of threshold. This effect was not due to run down, as no significant reduction was observed with vehicle controls (Figure 2B). In slices from GluA2-injected animals, we failed to reliably induce Ca^2+^ signals in PV+ interneuron dendrites, despite eliciting EPSCs of similar amplitude (Figure 2C). The reduction in the amplitude of Ca^2+^ responses was evidenced in both the mean ΔF/F and the signal-to-noise ratio. Significant differences between the responses to specific stimulation amplitudes were only detected at 300% threshold (Figure 2D). This difference remained significant even when Ca^2+^ signal amplitude was normalised to synaptic charge (t = 2.38, p = 0.023; unpaired t test), and there was no significant correlation between the Ca^2+^ signal amplitude at 300% stimulation and the corresponding evoked synaptic charge in either GluA2-injected or non-injected animals (Figure 2E). These data demonstrate that viral-driven expression of edited GluA2 successfully decreases CP-AMPAR expression and synaptically-evoked Ca^2+^ entry in hippocampal PV+ interneurons.

**Figure 2.**
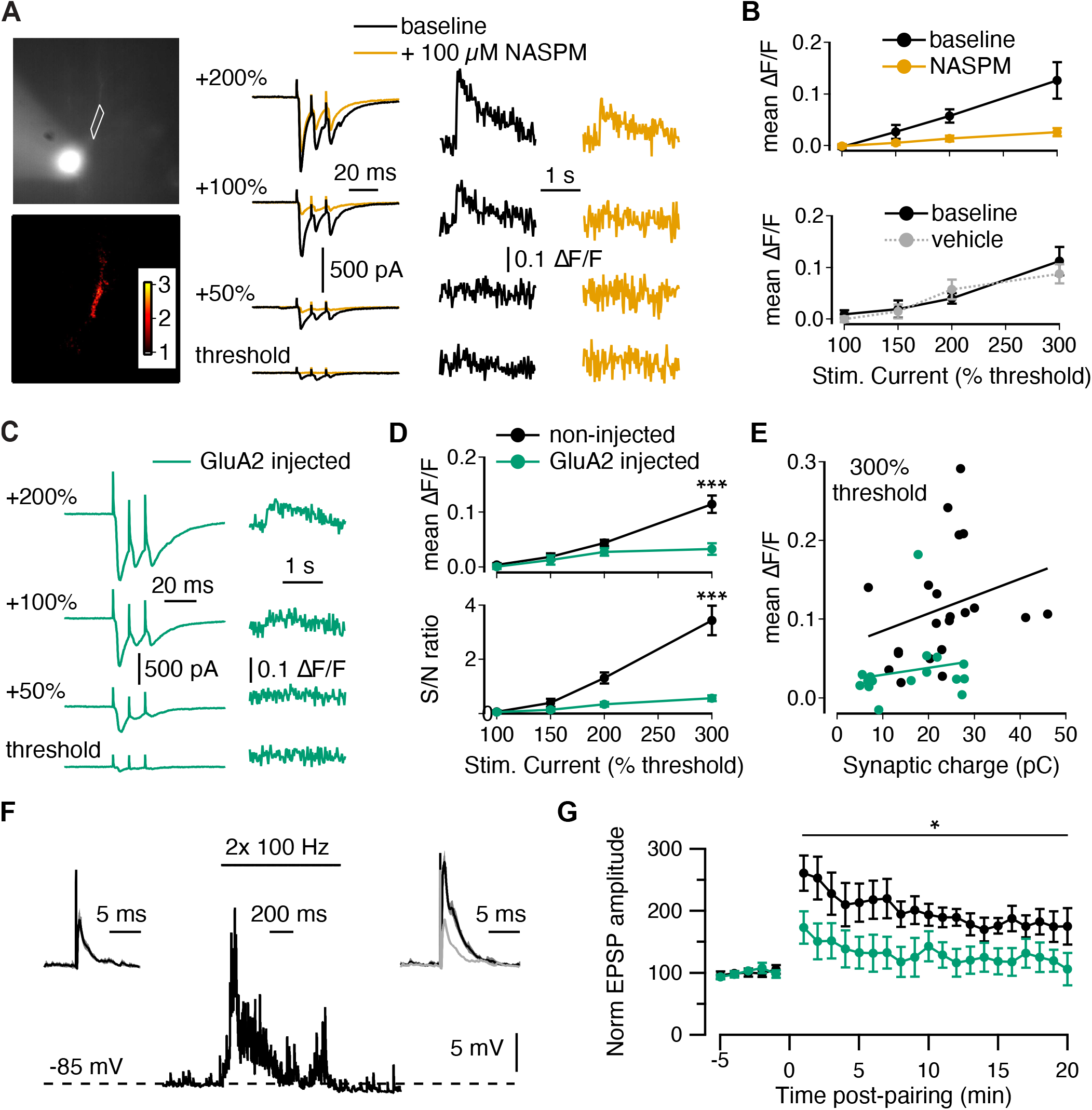
Viral transfection of PV+ interneurons with GluA2 reduces synaptically-evoked Ca_2+_ responses and the induction of anti-Hebbian plasticity. (A) Left: Image of PV+ interneuron loaded with 300 µM OGB-488, with ROI used for analysis (top). Template matching to the response in the ROI following 300% stimulation was used to calculate detection criteria (bottom) _45_, to confirm the localisation of the dendritic Ca_2+_ response. Right: EPSCs and dendritic Ca_2+_ signals recorded in response to synaptic stimulation (3 pulses at 100 Hz), before and after application 100 µM NASPM. (B) Changes in Ca^2+^ responses with increasing stimulation amplitude (F_(1.44, 8.62)_ = 16.13, p = 0.02) and application of NASPM (F_(1,6)_ = 8.07, p = 0.03 for main effect of drug; F_(1.25,7.48)_ = 7.59, p = 0.02 for interaction; repeated measures ANOVA), with significant effects observed at 200% (t = 3.71, p = 0.01) and 300% (t = 2.93, p = 0.03, n = 7; *post hoc* paired t tests) of threshold (top). Treatment with vehicle had no significant effects (F_(1,5)_ = 0.18, p = 0.69 for main effect of drug; F_(3,15)_ = 1.82, p = 0.19 for interaction; repeated measures ANOVA; bottom). (C) Representative EPSCs and dendritic Ca^2+^ signals recorded in PV+ neuron from GluA2-injected animal. (D) Comparison of dendritic Ca^2+^ responses in PV+ neurons from GluA2-injected animals and non-injected controls, as measured with either the mean ΔF/F (F_(1,35)_ = 10.46, p = 0.003 for main effect of group; F_(2.0, 77.6)_ = 8.3, p=0.001 for interaction; top) or the signal-to-noise (S/N) ratio (F_(1,35)_ = 11.04, p = 0.002 for main effect of group; F_(1.7, 57.8)_ = 10.7, p<0.001 for interaction; mixed ANOVAs; bottom). Significant differences between the responses to specific stimulation amplitudes were only detected at 300% threshold (mean ΔF/F: t = 3.96, p < 0.001; S/N ratio: t = 4.20, p < 0.001; n = 21 & 16 for control and injected, respectively; *post hoc* unpaired t tests). (E) Correlation between the Ca^2+^ signal amplitude at 300% stimulation and the corresponding evoked synaptic charge in PV+ neurons from either GluA2-injected animals (r = 0.18, p = 0.51) or non-injected controls (r = 0.28, p = 0.22; Pearson’s correlation test). (F) Sample traces from a PV+ cell from non-injected animal, showing excitatory postsynaptic potentials during baseline (left) and 20 minutes post-pairing (right; baseline response shown in grey). Pairing consisted of hyperpolarization and 2x 100 Hz stimulation for 1 s (middle). (G) Anti-Hebbian plasticity recorded in PV+ neurons from GluA2-injected animals non-injected controls (F_(1,16)_ = 6.06, p = 0.03 for main effect of group; F_(2.35, 37.54)_ = 0.45, p = 0.68 for interaction; n = 10 & 8 for control and injected, respectively; mixed ANOVA). *p < 0.05; ***p < 0.001. Values represent mean ± SEM.

To determine whether changes in CP-AMPAR expression in PV+ interneurons affect anti-Hebbian synaptic plasticity, we used a protocol in which cellular hyperpolarization was paired with 2x 100 Hz stimulation. This protocol reliably induced fast-onset, long-lasting potentiation of excitatory post-synaptic potentials (EPSPs) in PV+ interneurons from non-injected control mice but not in GluA2-injected mice (Figure 2F&G). Taken together, these results indicate that transfection of PV+ interneurons with our floxed GluA2 virus impairs both AMPAR-mediated Ca^2+^ influx and synaptic plasticity.

### Hippocampal PV+ interneuron transfection with the GluA2 virus causes a delay-dependent spatial working memory deficit

In order to investigate the effects of altering GluA2 AMPAR subunit expression in PV+ cells on behaviour, a cohort of PV-Cre/Ai9 mice received bilateral hippocampal (dorsal and ventral) injection of either the floxed GluA2 virus (n = 12; 6 males, 6 females) or a floxed GFP virus (control group; n = 11; 6 males, 5 females). The mice first performed a battery of tests to examine exploratory activity and anxiety, including the open field test, locomotor response to novelty in photocell cages, elevated plus maze, novelty preference Y-maze and food neophagia test (see Table S1 for results). The only notable differences in behaviour between the GluA2-injected and GFP-injected mice were a small but significant decrease in rearings in the locomotor test (p = 0.048), and a small non-significant increase in novelty preference in the Y-maze (p = 0.06). Here, we now focus on the effects on spatial learning and memory. PV+ interneurons in the hippocampus are necessary for optimal performance on spatial working memory tasks in rodents, such as the T-maze rewarded alternation (non-matching to place) task (Figure 3A)^14,18-20^. Mice were initially trained on the task (6 daily sessions of 10 trials) with a 5 s delay between the sample and choice phases of each trial. Overall, there was no difference between the performance of the two viral groups, nor any interaction between viral type and session (Figure 3B). However, while the performance of the control group improved across the trials within each session (first half vs second half of session; p < 0.001), the performance of the GluA2 group deteriorated (p=0.004), and the GluA2-injected mice performed significantly worse than controls during the second half of each session (Figure 3B), potentially reflecting proactive interference.

**Figure 3.**
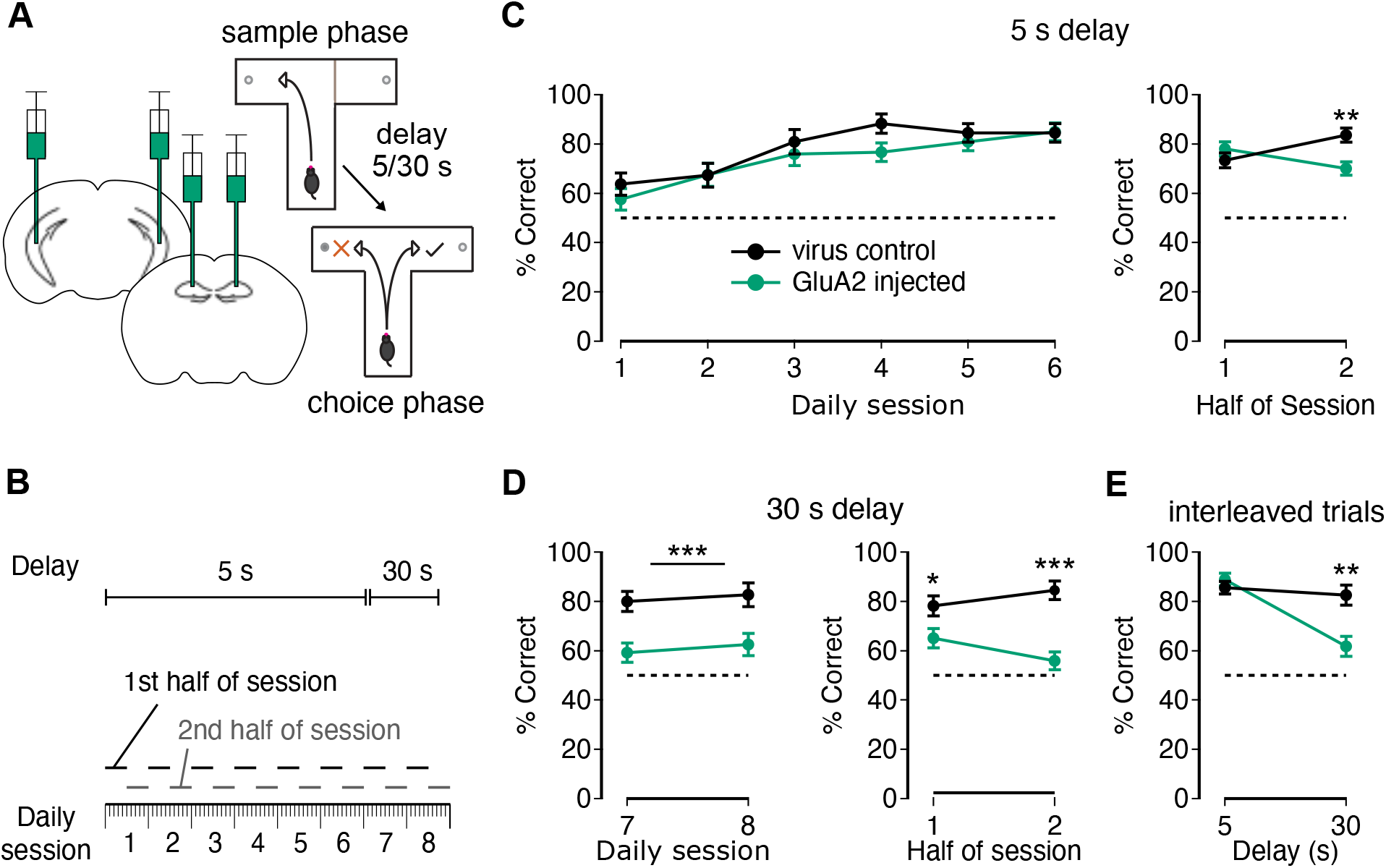
Overexpression of GluA2 in hippocampal PV+ interneurons leads to delay-dependent impairment in spatial working memory. (A) Illustration of viral injections across the extent of the hippocampus (left) and design of the T-maze task (right), with each trial consisting of a sample phase (top) and a choice phase (bottom) separated by a delay of 5 or 30 seconds. (B) Schematic of experimental design: each daily session consisted of 10 trials. For each delay, data were analysed across daily sessions and between the first (trials 1-5) and second (trials 6-10) halves of each session. (C) Left: Performance of GFP-injected controls (n = 11) and GluA2-injected group (n = 12) during training with a 5 s intra-trial delay (F_(1,21)_ = 1.42, p = 0.25 for main effect of viral treatment; F_(5,105)_ = 0.73, p = 0.60 for interaction; mixed ANOVA). Right: Change in performance across the first and second halves of each session (F_(1,21)_ = 25.78, p < 0.001 for interaction; First half: F_(1,21)_ = 1.32, p = 0.26; Second half: F_(1,21)_ = 11.67, p = 0.003; mixed ANOVA followed by *post hoc* simple main effects). (D) Left: Subsequent performance with a 30 s delay (F_(1,21)_ = 23.13, p < 0.001 for main effect of viral treatment; F_(1,21)_ = 1.05, p = 0.95 for interaction; mixed ANOVA). Right: Change in performance across the first and second halves of each session (F_(1,21)_ = 5.65, p = 0.027 for interaction; First half: F_(1,21)_ = 5.42, p = 0.03; Second half: F_(1,21)_ = 31.40, p < 0.001; mixed ANOVA followed by *post hoc* simple main effects). (E) Interleaving trials with delays of 5 and 30 s within the same session (F_(1,21)_ = 14.60, p = 0.001 for interaction; 5 s delay: F_(1,21)_ = 0.82, p = 0.38; 30 s delay: F_(1,21)_ = 13.91, p = 0.001 for effect of viral treatment; mixed ANOVA followed by *post hoc* simple main effects). *p < 0.04; **p < 0.008; ***p < 0.001. Values represent mean ± SEM.

Increasing the delay between the sample and choice phases within a trial can also have the effect of increasing proactive interference between trials^21^. When this intra-trial delay was extended to 30 s for two further sessions, the GluA2 group made significantly more incorrect choices compared to controls (Figure 3C). When these sessions were split into two halves as above, the GluA2-injected group now also performed significantly worse than controls during the first half of each session, with an even more pronounced difference for the second half of each session (Figure 3C). To rule out the possibility that this delay-dependent memory effect was due to order of testing, two further sessions were performed with interleaved delays of 5 or 30 s. The GluA2 group again made significantly more errors than controls with a 30 s delay, but not a 5 s delay (Figure 3D). These results are consistent with an increased effect of proactive interference in the GluA2 overexpressing mice.

### Expression of GluA2 in hippocampal PV+ interneurons impairs reversal learning in the Morris water maze

The mice were then tested on a hippocampus-dependent, appetitively motivated, spatial reference memory task on the elevated Y-maze^22^. Both groups acquired the task at an equivalent rate across the 10 sessions of testing and achieved the same high levels of performance by the end of training (Table S1)^22^. The animals were then assessed on the hippocampus-dependent, spatial reference memory version of the Morris water maze task (Figure 4A)^22^. Both groups acquired the task at a similar rate and reached the same level of asymptotic performance by the end of training. The GluA2-injected animals exhibited shorter latencies and path lengths to find the platform than controls during early training sessions, possibly due to them spending significantly less time near the side walls of the maze (Figure 4B)^23^. Probe tests during which the platform was removed from the pool showed that both groups remembered the designated platform location to the same extent (p = 0.44), with similar strong preferences for the ‘target’ quadrant (Figure 4C). These results further demonstrate the lack of a spatial reference memory impairment and no global hippocampal dysfunction among GluA2-injected mice.

**Figure 4.**
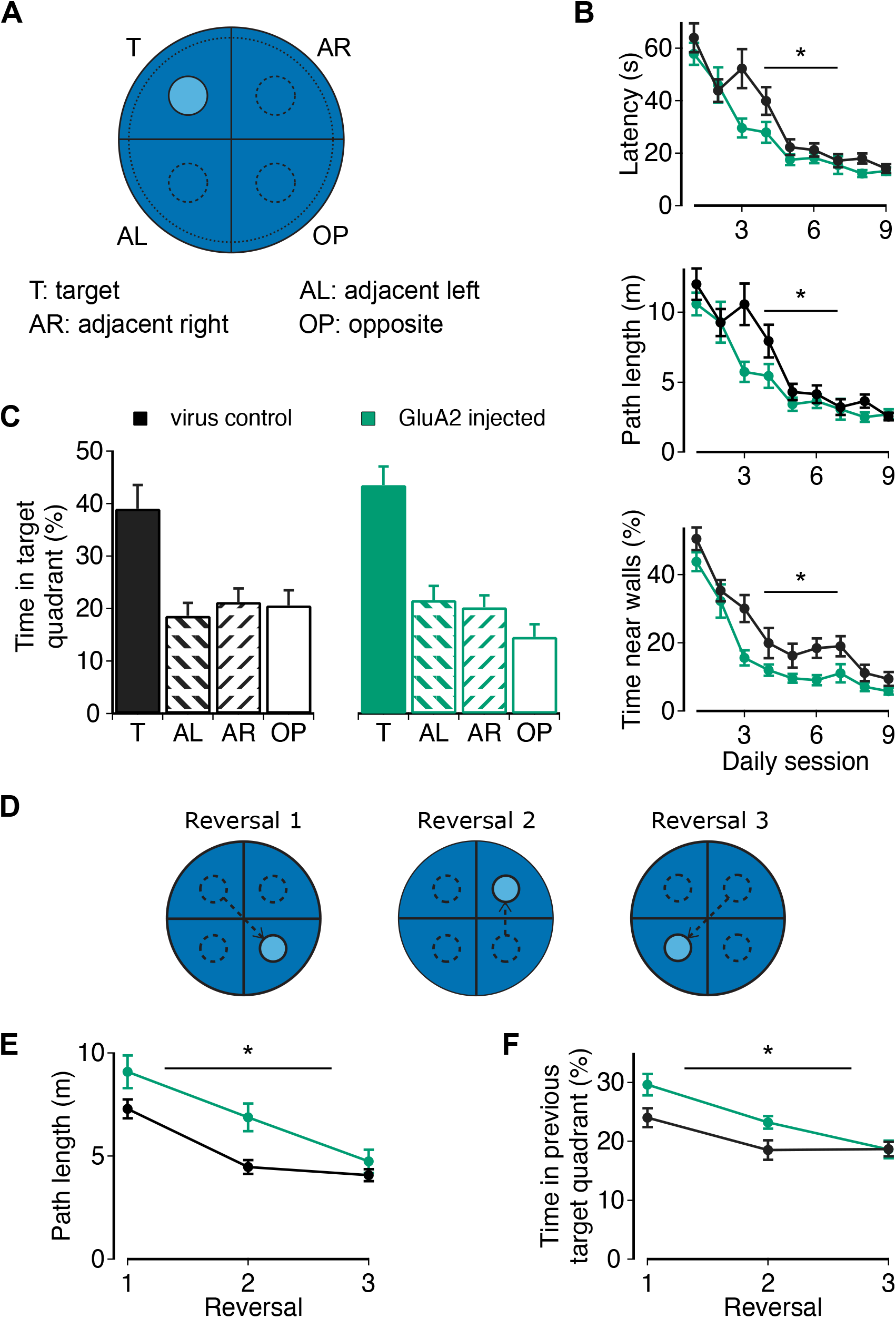
Overexpression of GluA2 in hippocampal PV+ interneurons impairs reversal learning without affecting acquisition of reference memory in the Morris water maze. (A) Illustration of water maze, including possible platform locations in the four quadrants of the pool. (B) Performance during training; latency (F_(1,21)_ = 6.29, p = 0.02; top), path length (F_(1,21)_ = 4.59, p = 0.04; middle) and thigmotaxis (F_(1,21)_ = 10.55, p = 0.004; bottom; main effects of viral treatment; mixed ANOVA). (C) Preference for the target quadrant averaged across both platform-free probe trials (t = 0.791, p = 0.44; unpaired t test). (D) Example sequence of changes in platform position during serial reversal phase of testing (platform sequences counterbalanced for viral treatment and gender). (E) Path length performance across all 3 reversals (F_(1,21)_ = 6.78, p = 0.02). (F) Time spent in the target quadrant from the previous reversal session (F_(1,21)_ = 5.82, p = 0.025; main effect of viral treatment; mixed ANOVA). *p < 0.05. Values represent mean ± SEM.

Once the mice had learned the original platform location, they were then subjected to a series of three spatial ‘reversals’, during which the platform was shifted to a new location in a different quadrant on each reversal (Figure 4D). Averaged across all three reversals, the GluA2-injected mice now exhibited significantly longer path lengths (Figure 4E) and spent a significantly greater percentage of time in the previous target quadrant (Figure 4F), indicating perseveration for the previously learned location. Overall, these data demonstrate that, whilst CP-AMPAR-mediated activity in hippocampal PV+ interneurons is not necessary for acquiring associative spatial memories, the loss of these receptors impairs the cognitive flexibility needed to disambiguate between competing memory traces. This indicates increased proactive interference, which would be consistent with findings from the T-maze spatial working memory task (Figure 3).

## Discussion

In this study we investigated the function of synaptic AMPAR-mediated Ca^2+^ influx in hippocampal PV+ interneurons by replacing CP-AMPAR with Ca^2+^-impermeable AMPARs, using cell type-specific viral expression of the Q/R-edited flop form of the GluA2 AMPAR subunit^24,25^. Similar approaches have been used to manipulate cellular AMPAR profiles in non-neuronal cell types^26,27^ and neurons^17,28-30^, but the behavioural effects of switching AMPAR profiles in hippocampal GABAergic interneurons remain unknown. Here we show that our viral approach yields sufficient expression of GluA2-containing AMPAR to (i) render carbachol-induced hippocampal gamma oscillations resistant to CP-AMPAR blockers, (ii) inhibit synaptically-evoked Ca^2+^ influx and anti-Hebbian plasticity in hippocampal PV+ interneurons, (iii) produce delay-dependent spatial working memory deficits which accumulated across trials within each session, and (iv) impair reversal learning on the Morris water maze, without deficits in initial acquisition of long-term spatial reference memory.

Previous studies have also reported selective spatial working memory deficits following manipulation of PV+ interneurons in mice^3,14,20,31^. In our present study, working memory deficits only emerged in GluA2-injected animals with both increasing numbers of trials within a session and with an increased delay, suggesting proactive interference during working memory retrieval. Likewise, although GluA2-injected animals exhibited normal acquisition of the Morris water maze, they did show a deficit in reversal learning, which is also consistent with increased proactive interference^32^. In the context of “winner-take-all” attractor network models, GluA2-injected mice disambiguate poorly between overlapping and competing hippocampal associative memory traces^33,34^. This suggests an important role for CP-AMPAR in the rapid and reversible recruitment of PV+ interneurons which, in turn, can inhibit the activity of the pyramidal cell ensembles that underlie these competing memory traces^1^. In this way, specific populations of PV+ interneurons may suppress the retrieval of inappropriate memories and thus reduce proactive interference. Given the importance of the prefrontal cortex in mitigating against interference in such memory tasks^35,36^, an intriguing possibility is that top-down signals from prefrontal cortex can instruct activity in the hippocampus to protect memory performance^37,38^. Our data demonstrate that this could involve the dynamic recruitment of PV+ interneurons, and notably the structure of hippocampal PV+ networks correlates with the propensity for reversal learning^39^.

Our data also suggest that the dynamic recruitment of PV+ interneurons involves anti-Hebbian plasticity via CP-AMPAR. We show that the overexpression of GluA2 in hippocampal PV+ interneurons reduced anti-Hebbian plasticity while maintaining synaptic transmission. Changing AMPAR subunit composition would be expected to alter synaptic kinetics^40^, which may alter cellular and circuit synchronisation. Indeed, we observed a small but significant reduction in the peak frequency of gamma oscillations, and overexpression of GluA2 across GAD67+ neurons has been shown to disrupt long-range gamma synchronisation^17^. However, GluA2-injected animals display cumulative and delay-dependent spatial working memory deficits, and show selective reversal learning deficits in spatial reference memory, which are more likely to reflect changes in plasticity rather than synaptic kinetics per se. In turn, the deficits in anti-Hebbian plasticity may shape gamma synchronisation^41,42^. Anti-Hebbian plasticity in PV+ interneurons may protect against memory interference by supporting the amplification of spatial selectivity in hippocampal assemblies^43^. The dynamic recruitment of PV+ interneurons through CP-AMPAR likely supports network oscillations and plasticity, which serves an important role in accurate memory retrieval and provides a promising clinical target for the treatment of memory deficits in Alzheimer’s disease and schizophrenia^44^.

## Acknowledgements

We thank the University of North Carolina Vector Core for custom preparation of the GluA2 viral construct. This work was supported by Wellcome Trust funding to the Oxford Ion Channel Initiative (M.C. and M.M.). M.A. is a PhD student in the NIH Oxford-Cambridge Scholars Program. This research was supported in part by the Intramural Research Program of the NIH.

## Author Contributions

M.C., D.M.B. and E.O.M. conceived the experiments. M.C. and M.M. performed the experiments. M.C. and M.M. analysed the data with support from D.M.B. and E.O.M. M.C., M.A., D.M.B. and E.O.M. wrote and edited the paper.

## Declaration of Interest

The authors declare no competing interests.

## STAR Methods

### RESOURCE AVAILABILITY

#### Lead Contact

Further information and requests for resources and reagents should be directed to and will be fulfilled by the Lead Contact, Ed Mann.

#### Materials Availability

All plasmids generated were sent to UNC Vector Core.

#### Data and Code Availability

The data and analysis code generated in this study are available upon reasonable request to the corresponding authors.

### EXPERIMENTAL MODEL AND SUBJECT DETAILS

All experiments were performed on adult mice of both sexes (6 weeks or older) homozygous for B6;129P2-*Pvalb*^*tm1(cre)Arbr*^/J (PV-Cre) and B6.Cg-*Gt(ROSA)26Sor*^*tm9(CAG-tdTomato)Hze*^/J (Ai9).

The animals were housed in holding rooms on a 12:12-hour light:dark cycle, with *ad libitum* food and water, except during the appetitively-motivated behavioural tasks, during which they were food-restricted to 85-90% of free-feeding body weight. Animals were allocated to experimental groups in a pseudorandom fashion. All testing was performed in the light phase.

All experiments were approved by the local ethical review committee at the University of Oxford and licensed by the UK Home Office.

## METHOD DETAILS

### Viral vectors and production

The *in vitro* experiments were performed in slices taken from either non-injected animals, or animals injected with a floxed GluA2 construct in an rAAV8 vector, whilst animals in the *in vivo* behavioural cohort were injected with either the GluA2 vector or an AAV8-CAG-FLEX-GFP reporter virus (Dr. Ed Boyden via UNC Vector Core). The virus was injected at 4.9 × 10^12^ PFU.

The GluA2 construct used in this study was generated with the assistance of GenScript and UNC Vector Core. A plasmid containing SalI and NotI restriction sites flanked by loxP sites downstream of an hSyn promotor was donated by Dennis Kaetzel (Ulm University) and sent to GenScript. Additionally, a sequence was designed encoding the ‘flop’ splice variant of mouse GluA2 cDNA, including an Q607R substitution that abolishes Ca^2+^ permeability, in an inverted sequence flanked by SalI and NotI sites. Between the signal peptide and GluA2 subunit, cDNA encoding an AU1 peptide tag was included, flanked by glycine-alanine linkers. This sequence was sent to GenScript to be synthesized, and restriction enzymes were used to transfer the sequences into the plasmid backbone between the loxP sites. This plasmid was sent to UNC Vector Core for amplification and packaging in an AAV8 serotype vector.

### Viral delivery and surgery

Mice were intrahippocampally injected with either the floxed GluA2 virus (*in vitro* and *in vivo* experiments) or the floxed GFP virus (*in vivo* experiments only).

Mice were anaesthetised in an induction chamber filled with 4% isoflurane and shaved on the scalp before being transferred to a stereotaxic frame, with anaesthesia maintained at a constant flow of 1-3% isoflurane in a 1.5 litres/minute oxygen flow, adjusted to maintain an approximately 1 Hz breathing rate. Systematic pain relief was provided with subcutaneous administration of 20 µl Buprenorphine (10 µg/ml) and 80 µl Meloxicam (250 µg/ml), with local pain relief delivered via administration of 20 µl (2.5 mg/ml) Bupivacaine under the scalp prior to incision. Following this, the skull surface was exposed via an incision of the scalp, and craniotomies were performed above the sites of injection.

Viral injections were made into the hippocampus. For histology, unilateral injections were made into either dorsal or ventral hippocampus. For *in vitro* experiments, bilateral injections were made into ventral hippocampus. For *in vivo* experiments, bilateral injections were made into both dorsal and ventral hippocampus. For all injections, 600 nl of virus was injected at each injection site over a period of 5 minutes, with the needle left in position for an additional minute following completion of injection before slow removal from the cranium. Dorsal hippocampal injections were made at ±2.1 mm mediolateral (ML), -1.5 mm anterioposterior (AP), and -1.5/-1.1 mm dorsoventral (DV), and ±3.0 mm ML, -2.8 mm AP, and -2.1/-1.5 mm DV. For initial histology, current-voltage relationship and field oscillation recording experiments, ventral hippocampal injections were made at ±2.75 mm ML, -2.7 mm AP, and -2.7/-3.1 mm DV, and ±2.9 mm ML, -2.9 mm AP, and -2.8/-3.2 mm DV. The latter coordinates were adjusted to ±3.25 mm ML, -3.0 mm AP, and -2.7/-3.1 mm DV for Ca^2+^ imaging, anti-Hebbian plasticity, and *in vivo* behavioural experiments.

After the completion of viral injection, the edges of the incision were treated with 20 µl (2.5mg/ml) Bupivicaine before the wound was sutured. Animals were provided 80 µl Meloxicam the following day for post-surgical pain relief, and were left for a minimum of 4 weeks to promote viral expression before experiments were performed.

### *In vitro* slice preparation

Adult mice were decapitated under isoflurance anaesthesia, and the brain removed from the skull and positioned on a Vibratome VT1200S (Leica) cutting stage whilst immersed in a 32°C sucrose-based cutting solution, containing (in mM): 150 sucrose, 15 glucose, 34.5 NaCl, 25 NaHCO_3_, 3 KCl, 1.25 NaH_2_PO_4_, 1 CaCl_2_, 10 MgSO_4_), with pH 7.2-7.4 and gassed with 5% CO_2_/95% O_2_ (carbogen). Horizontal sections of 350 µm thickness were transferred to a heated (∼31-32°C) interface recovery chamber filled with artificial cerebrospinal fluid (aCSF) of composition (in mM): 10 glucose, 126 NaCl, 24 NaHCO_3_, 3.5 KCl, 1.25 NaH_2_PO_4_, 2 CaCl_2_, 2 MgSO_4_, with pH 7.2-7.4 and gassed with carbogen. Slices were left in the recovery chamber for a minimum of 1 hour before any recordings were attempted.

### Local field potential recordings

Slices were transferred to an interface recording chamber and maintained at 34°C whilst perfused with aCSF plus 5 µM carbachol. An extracellular recording electrode (borosilicate, 1 MΩ) filled with aCSF was positioned in/near stratum pyramidale in area CA3 of ventral hippocampus. Oscillatory activity was recorded, and baseline activity was calculated by collecting frequency and power data at 1 hour following placement in the recording chamber.

Following this initial 1-hour recording period, recordings were ceased and restarted. After 5 minutes had been completed, either 100 µM NASPM or and equivalent volume of dH_2_O as control treatment was added to the perfusing solution, after which activity was recorded for 15 minutes. Once recordings were finished, the set-up was washed through with dH_2_O and then aCSF plus carbachol before further recordings were made.

### Recording EPSCs

Slices were transferred to a submerged chamber maintained at 31°C mounted on an upright microscope (Olympus BX51W1) and superfused with aCSF plus 50 µM DL-AP5 and 20 µM gabazine. Whole-cell recordings were made using recording electrodes (borosilicate, 4-7 MΩ) filled with a caesium methylsulfonate-based internal solution containing (in mM): 140 CsCH_3_SO_3_, 5 NaCl, 10 HEPES, 0.2 EGTA, 2 Mg-ATP, 2 Na-GTP, 5 QX-314, 1 spermine, 10 biocytin.

Recordings were made from PV+ interneurons and pyramidal cells in strata pyramidale and oriens (PV+ interneurons only) in area CA1 of ventral hippocampus, with cells recorded in whole-cell voltage-clamp mode. Series resistance was compensated throughout recordings (60-70%). AMPA currents were stimulated by the placement of a stimulating electrode (FHC) into stratum radiatum (for pyramidal cells) or stratum oriens (for PV+ interneurons). A stable current response size was established when cells were held at -70 mV by delivering a series of stimuli with a 10 s gap. Once a stable baseline was established, cells were subjected to a stimulation-step protocol, in which they received stimulations of constant amplitude every 15 s. Following 2 stimuli at -70 mV, the holding potential was raised to -60 mV; cells received 2 stimuli at each 10 mV step up to +80 mV. Cells that were still viable at +80 mV received a reversed stimulation-step protocol, after which cells were resealed, and slices were stored in 4% paraformaldehyde in 0.1 M phosphate buffer for histological cell identification.

### Ca^2+^ imaging in PV+ cell dendrites

As for current-voltage recordings, slices were transferred to a submerged chamber in an upright microscope and superfused with aCSF containing DL-AP5 and gabazine. A silver wire was connected to a stimulation box and placed inside a borosilicate pipette filled with aCSF (minus gabazine and DL-AP5). Cells were recorded in whole-cell voltage-clamp with borosilicate pipettes filled with the caesium methanesulfonate internal solution, plus 300 µM Oregon Green 488 Bapta-1 (OGB-488, Thermofisher Scientific). Cells were left for 15 minutes to allow OGB-488 to fill the cell, and the silver stimulation electrode was positioned next to a visible dendrite in the patched cell.

Cells were stimulated with increasing amplitude by the electrode until a visible current response (50-100 pA) was detected; this amplitude was determined the stimulation threshold. Subsequently, cells underwent a stimulation-recording protocol, during which they were stimulated 3 times at 100 Hz with increasing stimulation amplitude, starting at the stimulation threshold (100%), and then increasing to 150%, 200%, and 300% threshold. During stimulation, cells were visualised with 488 nm light and changes in fluorescence were recorded using MicroManager imaging software, with snapshots taken continuously at 50 Hz for 3 s (the stimulation was delivered 1 s into this window).

Following the completion of this protocol, the stimulation amplitude was returned to the threshold for current response induction, and adjusted accordingly if the threshold had shifted during the stimulation protocol. Cells were then treated with either 100 µM NASPM or an equivalent volume of control dH_2_O, and stimulated every 15 s for 10 minutes. Following this 10-minute period, the stimulation-imaging protocol was repeated. Once 300% threshold stimulation had been reached, recordings were finished from the cell and the recording electrode was withdrawn to reseal the cell membrane. The rig was washed out before further recordings were made.

### Anti-Hebbian plasticity in PV+ interneurons

As before, slices were transferred to a submerged chamber in an upright microscope and superfused with aCSF, this time without gabazine or DL-AP5. Whole-cell patch-clamp recordings were made in current-clamp mode, with borosilicate recording electrodes filled with a potassium gluconate internal solution containing (in mM): 110 KGluc, 40 HEPES, 2 ATP-Mg, 0.3 GTP-NaCl, 4 NaCl. PV+ interneurons in area CA1 of ventral hippocampus were recorded, with sections cut between area CA3 and CA1. Small excitatory post-synaptic potentials were evoked with the placement of a stimulating electrode in stratum oriens. Following break-in, cells were injected with slow current to maintain a stable potential of - 70 mV, and stimuli were delivered every 15 s. Once a stable baseline of sub-10 mV evoked potentials was established over a 5-minute period, current was injected to hyperpolarise the cell to between -85 and -90 mV, and two rounds of 1 s 100 Hz burst stimulation were delivered before the cell was returned to the previously-held potential. Following this, stimuli were delivered every 15 s as before, with the same stimulation amplitude, for 20 minutes. After 20 minutes, the recording was terminated, and the cell subjected to a simple step protocol (10 steps, increasing in steps of ±20 pA) before recordings were completed.

### Behavioural tasks

All behavioural tasks were performed in the same cohort of 24 mice; 11 injected with the floxed GFP control virus (6 males, 6 females), and 12 injected with the floxed GluA2 virus (6 males, 6 females). One female control animal had an abnormal hyperactive phenotype on the locomotor assay and was thus excluded from the analysis. Behavioural tests were performed in the following order: elevated plus maze, locomotor activity assay, open field test, Y-maze spatial novelty preference, food neophagia, T-maze rewarded alternation spatial working memory task, elevated Y-maze spatial reference memory task, water maze spatial reference memory task and spatial reversals. All tasks were performed in separate experimental rooms.

#### Elevated plus maze

A plus-shaped maze elevated 70 cm above the floor consisted of four arms of equal length (35 × 6 cm) radiating from a central square (6 × 6 cm) at right angles. Two opposing arms (the ‘closed’ arms) were enclosed in a perimeter of 20 cm high walls, with two opposing ‘open’ arms placed between the closed arms. The maze was placed in a quiet, dimly-lit room (15 lux light intensity on the open arms, 9 lux in the centre square). Each animal was placed on the maze in the centre square and allowed to explore the maze for 5 minutes. All animals in the cohort were run individually on a single day, with all males run on the task before females. Animals were moved into the testing room in their home cages a minimum of 5 minutes before testing.

#### Locomotor assay

Spontaneous locomotor activity (LMA) was tested in a novel environment, during which animals were placed individually in transparent Perspex cages (26 cm x 16 cm x 17 cm) surrounded by two photobeam perimeter frames (PAS-Open Field, San Diego Instruments) stacked on top of each other. Horizontal movements within the cage disrupt the photobeams between opposing frames, and accumulation of beam breaks represents activity levels. Animal rearing results in beam breaks in the upper frame, enabling distinction between these two types of activity. Animals were run in sub-cohorts of 6 mice (plus a 5) on successive days at the same time of day (1 pm start time), with the cohort counterbalanced such that at least one animal of each gender and viral treatment was tested in each of the cages. Animals were placed in the cages for 2 hours before being returned to their home cages; horizontal and vertical beam breaks were separated into 5-minute time bins for analysis.

#### Open field test

A 60 cm diameter, white metallic cylindrical arena with high walls was brightly illuminated from above. Ethovision XT split the arena into 3 sections; a 10 cm radius inner circle, an outer circle of 15 cm radius, and a perimeter region of 5 cm radius. Prior to testing, animals were housed individually in cages for 5 minutes, then moved into the testing room and immediately placed inside the arena at the edge of the perimeter wall. Testing lasted for 5 minutes, at which point mice were removed and rehoused with their cage mates. As with the elevated plus maze, all animals were tested on the same day.

#### Spatial novelty preference

A clear Perspex Y-shaped maze with 30 cm x 8 cm x 20 cm arms was placed in a room with multiple extramaze spatial cues, with the floor of the maze lined with sawdust. Mice were assigned a ‘start’ arm, a ‘to-be familiar (other) arm’ and a ‘novel arm’, counterbalanced across the cohort for gender and viral treatment. Animals were moved into individual cages 5 minutes before testing on the maze. They were then moved into the testing room and immediately placed into the ‘start’ arm. During the exposure phase of the trial, only two arms were accessible, the ‘start’ and the ‘other’ arm; the ‘novel’ arm was blocked off with a black plastic block. The animal explored the two available arms for 5 minutes starting from the time it first left the ‘start’ arm. After 5 minutes, the animal was removed and returned to its temporary holding cage for 1 minute, during which the black block was removed and the sawdust brushed around the maze to diminish any olfactory clues. The animal was then returned to the start arm for the test phase, which lasted for 2 minutes, again starting from when the animal first left the start arm. The mouse was now free to explore all three arms. The animal was then returned to its home cage. All animals were tested on the same day according to a counterbalanced sequence.

#### Food neophagia

Prior to testing, animals were food restricted overnight for approximately 16 hr. Animals were transported into the testing room a minimum of 5 minutes before testing. During a trial, mice were placed underneath an upturned translucent jug of 15 cm diameter, with a 2 cm spout. Beneath the spout, a food well was filled with a 50:50 condensed milk:water solution that the animals had not previously encountered. The time until first contact with the milk and first consumption were recorded. Animals were left for 2 minutes, or until they first consumed the milk. Animals that did not drink from the well within 2 minutes were removed from the jug and held in a separate cage for approximately 2 minutes, before they were then replaced under the jug for another period of maximum 2 minutes or until they began drinking. For these mice, latency to first consumption was summed across both test periods (i.e. max latency was 240 s).

#### Rewarded alternation T-maze task

Spatial working memory was tested using an elevated, wooden T-maze with a start arm (47 cm x 10 cm), two goal arms (35 cm x 10 cm), and a 10 cm high perimeter wall. Plastic sliding doors were used to selectively block entry to goal arms, and plastic food wells were located at the end of each goal arm. The maze was placed in a dimmed room with distinct extramaze spatial cues. The 50:50 milk:water solution was used as a reward throughout testing.

Food-deprived mice were first habituated to the maze. Animals received several 10-minute habituation sessions on the maze, first with cage mates and then individually, during which they explored the maze and received rewards in the food wells at the end of the goal arms. Following extensive habituation, all animals were running well on the maze and readily consuming the milk rewards.

During the first phase of the experiment, animals received 6 sessions comprising 10 trials. A trial involved two phases: a sample phase and a choice phase. During the sample phase, one arm was blocked by a plastic door, so when the animal was placed in the start arm, it was forced to enter the open goal arm and could then consume the milk reward in the food well. After this, the animal was removed from the maze for 5 s, during which the previously blocked arm was opened by removing the block, and any trace of milk in the sampled food well was removed. The animal was then placed back in the start arm for the choice phase of the trial, during which the animal had to enter the previously blocked arm to receive a second reward (i.e. it was rewarded for alternating). Entering this arm was considered a correct choice, whilst if it entered the previously visited goal arm, this was considered an incorrect choice and the animal was removed from the maze without receiving a second reward. A choice was considered to be made when the animal placed 4 paws inside one of the goal arms. The animal was then removed from the maze for 2 minutes before receiving the next trial; animals were run in pairs, so each animal would perform a trial during the inter-trial interval of the other mouse. During each session, the right and left goal arms were used as the sample arm for 5 trials each, presented in a pseudorandom order with no more than 3 consecutive trials having the same sampled goal arm.

After these 6 sessions, mice received 2 more testing sessions of 10 trials; however, the intra-trial delay between the sample and choice phases was now increased from 5 s to 30 s. After these two days, the mice had 2 final sessions consisting of 12 trials presented in a pseudorandom order, 6 with the 5 s delay and 6 with the 30 s delay. There were 3 trials with the right arm as the sampled arm for each delay and 3 trials with the left arm as the sampled arm; there were no more than 3 consecutive trials during each session with either the same delay or sampled arm.

#### Elevated Y-maze spatial reference memory task

A Y-shaped maze with 3 arms (50 cm x 9 cm, with 0.5 cm high walls) radiating out from a central hexagon (14 cm diameter) at 120° angles was elevated 80 cm above the ground on a detachable base upon which the maze could be rotated. Food wells were placed 5 cm from the far end of each arm. Animals received one day of habituation to the maze in a separate room prior to training. The maze was then located in a well light room with several extramaze spatial cues. Mice received 10 sessions of testing, with 10 trials per day. Each animal was assigned a reward arm defined by the allocentric, extramaze spatial cues (counterbalanced for viral treatment and gender). Mice received 5 trials per session with each of the other two arms as the start arm, in a pseudorandom order with no more than 3 consecutive trials from the same start arm. A trial began with a mouse being placed at the far end of the start arm, and proceeded until the animal entered the correct arm and consumed the reward, or entered the incorrect arm and was removed from the maze. Trial latency and choice accuracy were recorded for each trial. The maze was rotated approximately every 2-4 trials to ensure the animals navigated using extramaze cues instead of intramaze cues. Animals were held in individual cages during testing, and were run in groups of 4, with a 5-minute inter-trial interval for each mouse. On the tenth day, to control for the possibility that the mice were using the smell of the reward to choose the correct arm, the reward was only delivered to the well after the animal had correctly entered that arm (post-choice baiting).

A majority of the cohort initially perseverated in making the same egocentric body-turn, independent of the start arm, resulting in 50% correct choices per session. In order to counteract this persistent tendency, after completing session 5, all animals scoring 80% or lower on that day were given ‘remedial training’, during which they were given blocks of trials (minimum 1-minute inter-trial interval) starting from the arm they consistently made the incorrect decision from. They would receive these trials until they consistently changed their decision-making to enter the reward arm (3 correct trials). After completing this, they then received a block of trials from the other start arm to ensure they hadn’t simply altered their turning response. After 2 correct trials from this arm, animals were returned to their home cages. This process was repeated after session 6 for any animal scoring 60% or lower (n=7 animals). No further remedial training was provided after session 6.

#### Water maze task

Spatial reference memory was assessed using a 2 m diameter, 0.9 m tall circular watermaze elevated 36 cm above the ground, in which a 21 cm diameter circular platform could be placed. The pool was filled with water to a depth of 1 cm above the surface of the platform, and was turned opaque with 600 ml white ready-mixed paint (Hobbycraft). The platform could be positioned in one of 4 possible locations, assigned northwest (NW), northwest (NE), southeast (SE) and southwest (SW), corresponding to the quadrant of the pool. The platform positions were 50 cm from the edge of the pool and equidistant from each other and the centre of the maze.

#### Acquisition

Across 9 days of acquisition training, animals received 4 trials per day, starting from one of 8 possible positions at the perimeter of the pool (NW, N, NE, E, SE, S, SW, W), in a pseudorandom order. Animals were assigned a fixed platform location throughout acquisition training, in either the NW or SE quadrants, counterbalanced by viral treatment and gender. The animal was allowed to swim for either 90 s, or until it found the hidden platform. In the former case, after 90 s the mouse was guided to the platform location. Once on the platform, the animals were left for 30 s, before they were removed, dried with a towel, and placed back in the pool for their next trial. WaterMaze software (Actimetrics) was used to track the animal’s position. Various parameters, including trial latency, path length, swimming speed, time spent at side walls, time spent in each of the four quadrants of the maze, etc., were recorded by the software for analysis.

#### Probe tests

The day after sessions 6 and 9 of testing, the animals received a 60 sec probe test, during which the platform was removed from the maze and the animal was placed into the water at a starting point opposite the previously learned platform location. Mice were allowed to swim freely for 60 sec and the time spent in each of the 4 quadrants of the pool was recorded. After watermaze testing, animals were moved to a 31°C incubator to dry before returning to their home cage.

#### Reversal testing

Following acquisition of this reference memory task, the mice underwent reversal testing, consisting of three rounds of training in which the platform location was moved to a novel location in the watermaze. During the first round, the platform was moved to the quadrant opposite its previous location for each animal (e.g. platform moved from NW to SE; Figure 3D). The animals had 3 days of testing (with 4 trials per day) to learn this new location, followed by a probe test at the end this training. During the second round, the platform location was moved again to another new location in either the SW or NE quadrants, with these new locations assigned to the mice in a way that was counterbalanced for viral treatment, gender, and previous platform position. Again, animals received 3 days of training, followed by a probe test. Finally, the platform location was moved to the opposite quadrant (NE or SW; i.e. the last remaining unused quadrant), and mice received 3 more days of testing plus a final probe test. Control mice failed to show any significant quadrant preference during these probe tests and so these data are not reported.

### Histology

Animals from the *in vivo* behavioural cohort, and those used for full-brain histology, were perfused, and their brains extracted. Animals received a 0.4 ml (200 mg/ml) intraperitoneal injection of pentobarbital (Euthatal, Merial Animal Health Ltd.), before being transcardially perfused with first phosphate buffered saline (PBS), followed by paraformaldehyde (PFA, 4% in 0.1 M phosphate buffer, Taab). Fixed brains were extracted from the skull and stored at 4°C in PFA. Brains were sectioned on a VT1000S vibratome (Leica), with horizontal (for ventral hippocampus) or coronal (for dorsal hippocampus) sections cut with 60 um thickness. Sections were cut with PBS, and washed 3 times before spending 20 minutes in 0.15% tris-buffered saline-Triton X-100 (TBS-T). Sections then spent 1 hour in 20% normal horse serum (NHS) in TBS-T, briefly washed in TBS-T, and then incubated for 48 hours at 4°C with the primary antibody (mouse anti-AU1, BioLegend) diluted 1:1000 in 2.5% NHS in TBS-T. After this incubation, slices were washed thrice in TBS-T before overnight incubation at 4°C with the secondary antibody (Alexa Fluor-488-coupled goat-anti-mouse, ThermoFisher Scientific) diluted 1:500 in 2.5% NHS in TBS-T. Finally, sections were washed three times in PBS, once in phosphate buffer, and mounted with Fluoroshield (Sigma Aldrich). All sectioning, incubations, and washes not done at 4°C were done at room temperature. Individual slices in which single-cell voltage-clamp recordings were made were stored in 4% PFA at the end of recordings and subsequently stained for AU1 expression and biocytin. The slices underwent the same histology procedure as above, except in addition to overnight incubation with the secondary antibody, sections were also incubated with AMCA Avidin D (Vector Labs) diluted 1:1000 into the same solution as the secondary antibody. They were then mounted the following day as above.

Mounted sections were imaged at 5x, 20x, and 40x using a Leitz DRMD Fluorescence Microscope (Leica) and a Retiga 2000R CCD camera (QImaging). Images of fluorescence were taken using Ocular software. Additional images were taken using a FV1200 Confocal Microscope (Olympus) fitted with PMT detectors, using FV10-ASW 4.2 Fluoview software, at 40x magnification.

## QUANTIFICATION AND STATISTICAL ANALYSIS

### Network oscillations

Effects of viral treatment and drug application on oscillation properties were determined by studying the peak frequency and power area (20-80 Hz) of recorded field potentials. Recorded data was binned in 5 s sweeps. Peak frequency and power area values were calculated for each sweep, and baseline values for each slice were determined by taking the mean of 12 consecutive sweeps at the 1-hour mark. Minute-by-minute values during the pharmacological experiments were calculated by taking the mean value of 5 consecutive sweeps around each minute mark. Power area values were normalised across slices via logarithmic conversion.

Differences in baseline oscillation power and frequency properties between non-injected slices and slices taken from GluA2 virus-transfected animals were determined using unpaired t tests. Changes in power and frequency following vehicle or NASPM solution treatment were calculated between baseline and 10 minute post-application, and analysed using 2 (treatment) * 2 (slice type) ANOVA, with any interactions explored with *post hoc* unpaired t tests.

### Rectification index

Rectification properties of cells were determined by calculating the amplitude of the second response at each voltage step. The rectification index was calculated as the amplitude of the current response 40 mV above the reversal potential compared to the response at 40 mV below the reversal potential. Cells were excluded from analysis if, during the stimulation-step protocol, the series resistance went above 30 MΩ or varied by >15% within the rectification index window (±40 mV of the reversal potential). For cells that it was possible to perform two stimulation-step protocols in, the one with the lower variation in series resistance was used for analysis purposes. Due to the heterogeneous distribution of the rectification index, particularly within the virus-treated PV+ interneuron group, non-parametric Mann-Whitney U tests were used to test for differences between the control and virus-treated PV+ interneuron populations. The remaining comparisons were performed using unpaired t tests. Using either parametric or non-parametric tests for all comparisons did not alter the pattern of significant results.

### Spontaneous EPSCs

Spontaneous EPSCs were detected during the baseline stimulation period for experiments examining rectification, when the cells where voltage-clamped at -70 mV. Events were initially detected as those points exceeding 5 pA above a running 2-ms baseline mean, and events less than 5 s.d. above the baseline noise rejected. To avoid multiple detections of large events, the event detection algorithm recommenced after the peak of each detected event. Event detection was checked by visual inspection. Properties including current amplitude, area, and decay rate were calculated from the mean waveforms. The median for each of these variables across the baseline period was determined for PV+ interneurons in the control and GluA2 virus-transfected groups. Due to the heterogeneous distribution of IEIs, particularly within the virus-treated PV+ interneuron group, non-parametric Mann-Whitney U tests were used to test for differences between the control and virus-treated PV+ interneuron populations. The remaining comparisons were performed using unpaired t tests. Correlations between rectification indices of cells and spontaneous current properties were determined for each PV+ interneuron group by calculating bivariate Pearson’s correlation coefficients. Using either parametric or non-parametric tests for all comparisons did not alter the pattern of significant results.

### Dendritic Ca^2+^ signals

Ca^2+^ signals in PV+ dendrites were quantified within manually-defined regions of interest (ROI). The ROI response at 300% threshold stimulation was used in a template matching algorithm ^45^ to calculate detection criteria, and confirm the localisation of the dendritic Ca^2+^ response, and the ROI updated as necessary. The extracted Ca^2+^ transients were bleach-corrected by fitting an exponential curve function through both the baseline and end of the recording window. The signals were quantified as the mean ΔF/F for the 2 s period following stimulus onset, with signal-to-noise ration calculated by normalizing by the baseline standard deviation. The effects of NASPM or control treatment on signal amplitude compared to pre-treatment baseline were determined using 2 (drug treatment) *4 (stimulation amplitude) repeated measures ANOVA, with interactions explored using *post hoc* paired t tests. The effects of viral treatment were determined using 2 (viral treatment) * 4 (stimulation amplitude) mixed ANOVAs, with interactions explored using *post hoc* unpaired t tests. Relationships between current area and Ca^2+^ signal amplitude following 300% threshold stimulation were determined for each PV+ interneuron group by calculating bivariate Pearson’s correlation coefficients.

### Synaptic plasticity

Cells in which the input resistance drifted by 20% or more throughout the 25 minutes of baseline and post-pairing stimulation were excluded from analysis. The amplitudes of evoked potentials were determined, and means were calculated for 1-minute time bins. EPSP amplitudes were normalised to the mean amplitude of evoked EPSPs during the 5-minute baseline period. Significant differences in post-pairing EPSP amplitudes between the control and GluA2-treated cell populations were determined using a mixed ANOVA.

### Analysis of behaviour

#### Elevated plus maze

Ethovision XT (Noldus) was used to track the animal’s movement in the maze and calculate time spent in, latency to enter, and number of entries into the open and closed arms, as well as total distance travelled. Differences between the two viral groups were determined using between-subjects, unpaired t-tests.

#### Locomotor assay

Differences in overall locomotor activity levels and rearing between the two viral groups were determined using unpaired t tests, whilst differences in activity across time were determined using a 2 (viral types) * 24 (time bin) mixed ANOVA; any interactions were explored *post hoc* with simple main effects.

#### Open field test

Ethovision tracked time spent, entries into, and latency to first entry into the outer and inner circles, as well as the perimeter region of the open field arena. Differences between groups were determined using unpaired t tests.

#### Spatial novelty preference

During both phases, arm entries and time spent in each arm were manually recorded by the experimenter, with an arm entry defined as the animal having all four paws inside an arm, and an arm exit defined as the animal removing all four paws from the arm. A novelty preference ratio was calculated during the test phase for the time spent in the ‘novel’ arm compared to the ‘other’ arm, using the formula [novel arm/(novel + other arm)]. Differences in entries into the ‘novel’ arm and in the novelty preference ratio between the two viral groups were determined using unpaired t tests.

#### Food neophagia

The latency to first contact and to first consumption of the milk from the food well were recorded, and differences between the two groups assessed using unpaired t tests.

#### Elevated Y-maze spatial reference memory task

Differences in performance between the two groups across the training/testing period were determined using a 2 (viral type) * 10 (session) mixed ANOVA.

#### Rewarded alternation T-maze task

For each trial, the latency to sample, the choice latency, and the decision (correct or incorrect) was recorded. Differences in performance on the task between the two groups during the 5 s delay testing period were assessed using a 6 (session) * 2 (viral type) mixed ANOVA. Performance across the two days with a 30 s delay was averaged across the two sessions for each animal, and differences between the two groups were determined using an unpaired t test. Performance across the two days with interleaved short/long-delay trials was averaged across the two sessions, and delay-dependent differences between the two groups ascertained using a 2 (delay) * 2 (viral type) mixed ANOVA, with any significant interactions explored *post hoc* using analysis of simple main effects. Differences in performance over the course of sessions were explored by splitting sessions into 2 blocks of 5 trials (first half; trials 1-5 and second half; trials 6-10) and compared using mixed ANOVAs.

#### Water maze task

Differences in trial latency, path length and % time spent near the maze perimeter (thigmotaxis) during watermaze training between the two viral treatment groups were tested using 9 (session) * 2 (viral type) mixed ANOVAs. Times in target quadrant during the probe tests were averaged across both probe sessions, and differences determined using unpaired t tests. Path length and time spent in the previous target quadrant were averaged across the multiple spatial reversals and differences between the viral treatment groups were determined using mixed ANOVAs.

### Statistics

Statistical analysis was performed in SPSS. Significance values for all two-tailed statistical tests were set at 0.05.

## KEY RESOURCES TABLE

| REAGENT or RESOURCE | SOURCE | IDENTIFIER |
| --- | --- | --- |
| **Antibodies** |  |  |
| mouse anti-AU1 | BioLegend | Cat# 901901, RRID:AB_2565013 |
| AlexaFluor 488 goat anti-mouse | Thermo Fisher Scientific | Cat# A-11001, RRID:AB_2534069 |
| **Bacterial and Virus Strains** |  |  |
| AAV8-CAG-FLEX-GFP | UNC Vector Core | N/A |
| Floxed GluA2 | Custom Prep (UNC Vector Core) | N/A |
| GluA2 plasmid | Custom Prep (GenScript) |  |
| **Chemicals, Peptides, and Recombinant Proteins** |  |  |
| NASPM | Tocris | Cat# 2766 |
| DL-AP5 | Tocris | Cat# 0105 |
| Gabazine | Abcam | Cat# ab120042 |
| Oregon Green 488 Bapta-1 | Thermo Fisher Scientific | Cat# O6806 |
| **Experimental Models: Organisms/Strains** |  |  |
| B6;129P2-*Pvalb*<sup>tm1(cre)Arbr</sup>/J | The Jackson Laboratory | Stock# 008069, RRID:IMSR_JAX:017320 |
| B6.Cg-*Gt(ROSA)26Sor*<sup>tm9(CAG-tdTomato)Hze</sup>/J | The Jackson Laboratory | Stock# 007909, RRID:IMSR_JAX:007909 |
| **Software and Algorithms** |  |  |
| Micro-Manager | NIH | RRID: SCR_000415 |
| ImageJ | NIH | RRID: SCR_003070 |
| IGOR Pro | Wavemetrics | RRID:SCR_000325 |
| SPSS | IBM | RRID:SCR_002865 |
| SnapGene Viewer | SnapGene | RRID:SCR_015053 |
| Ocular software | QImaging | http://www.qimaging.com |
| Fluoview | Olympus | RRID:SCR_014215 |
| Ethovision XT | Noldus | RRID:SCR_000441) |
| WaterMaze | Actimetrics | http://www.actimetrics.com |
| PAS-Open Field | San Diego Instruments | http://www.sandiegoinstruments.com |

## Notes

### Competing Interest Statement

The authors have declared no competing interest.

