## Supplementary Material for "Calcium-permeable AMPAR in hippocampal parvalbumin-expressing interneurons protect against memory interference"

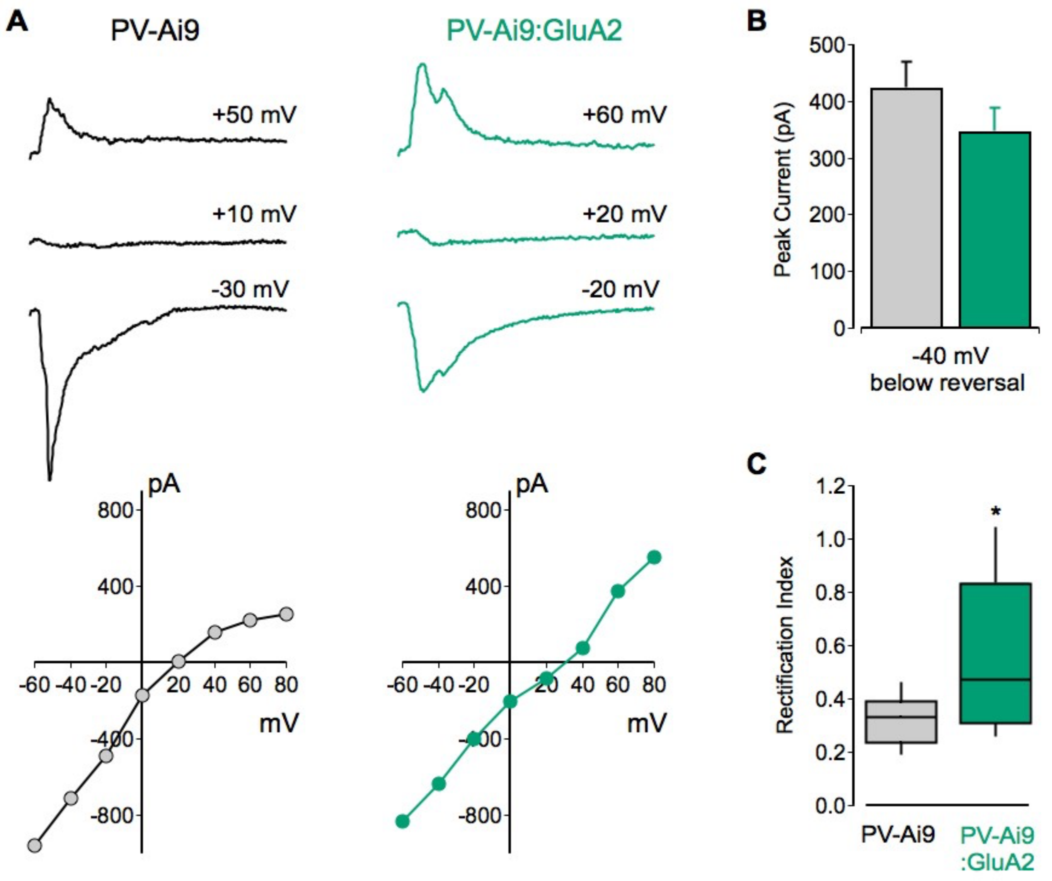

**Figure S1. Over-expression of GluA2 reduces inward rectification of evoked EPSCs in PV+ interneurons.**

(A) Example traces of evoked EPSCs recorded in PV+ interneurons at +/-40 mV from the reversal potential (top), and corresponding I-V plots (bottom), in slices prepared form non-injected (left) and GluA2-injected mice (right).

(B) There were no significant differences in the peak of evoked ESPCs recorded at -40 mV below the reversal potential ($p = 0.27$; unpaired t test; $n = 12$ for each group). Values represent mean ± SEM.

(C) The rectification index was significantly increased in PV+ cells from GluA2-injected mice compared to non-injected controls ($U = 115$, $p = 0.01$; Mann-Whitney U test). Box-and-whisker plot shows the median, where the box represents interquartile percentile range, and the whiskers represent the $10^{th}$ and $90^{th}$ percentiles.

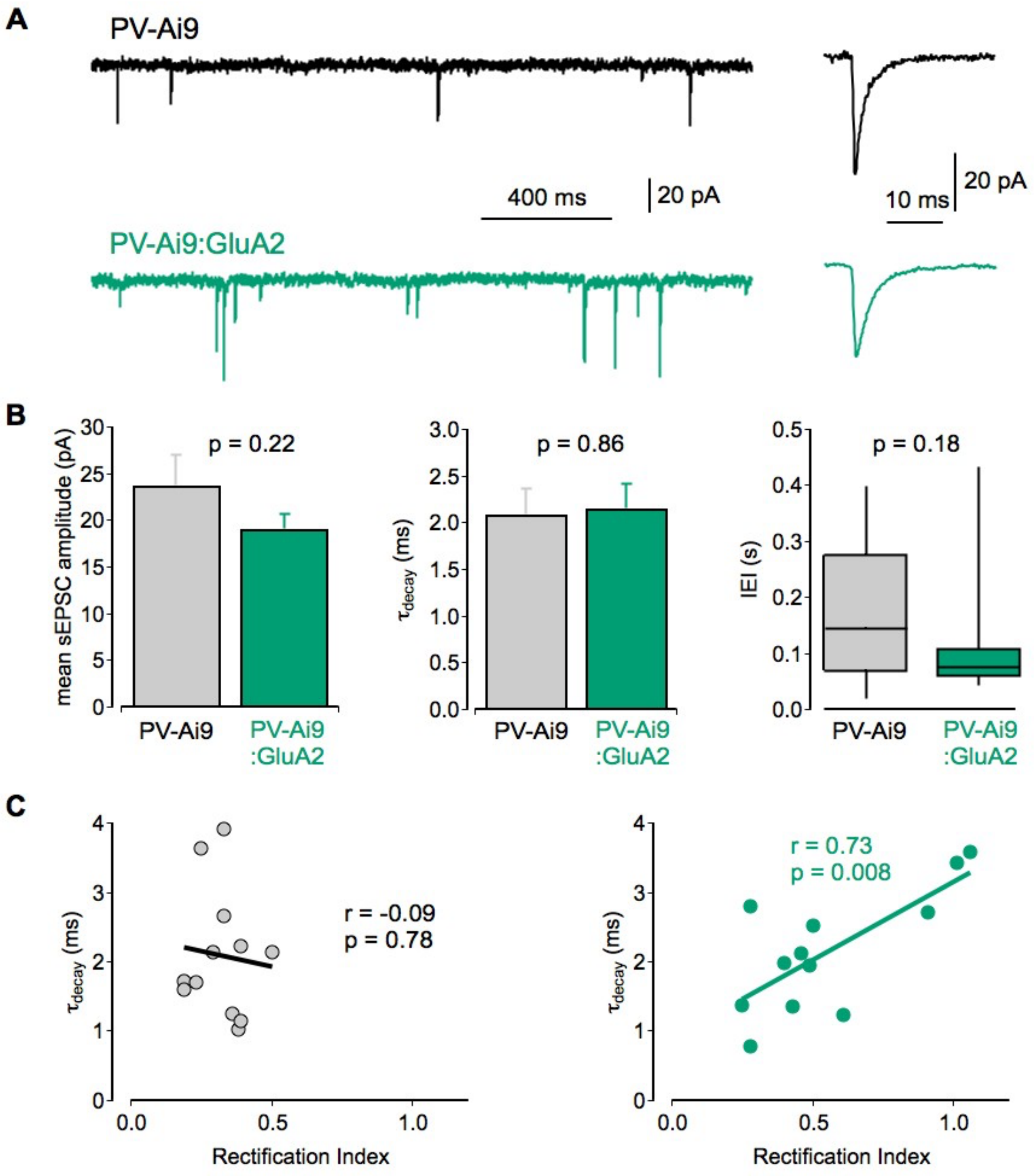

**Figure S2. Effect of GluA2 over-expression on spontaneous EPSCs in PV+ interneurons**

(A) Example traces of sEPSCs recorded in PV+ interneurons at -70 mV (left), and the average waveforms (right), in slices prepared form non-injected (top) and GluA2-injected mice (bottom).

(B) Over-expression of GluA2 in PV+ interneurons had no significant effect on mean sEPSC amplitude (left; $t = 1.27$, $p = 0.22$; unpaired t test), time constant of decay (middle; $t = 0.15$, $p = 0.86$; unpaired t test) or inter-event interval (IEI) (right; $U = 48$, $p = 0.18$; Mann-Whitney U test; $n = 12$ for each group).

(C) Plots of time constant of decay of sEPSCs against rectification index of evoked EPSCs, showing a significant positive correlation in PV+ cells from GluA2-injected mice (Pearson's linear correlation). Values represent mean ± SEM, except for box-and-whisker plot that shows the median, where the box represents interquartile percentile range, and the whiskers represent the 10$^{th}$ and 90$^{th}$ percentiles.

| Behavioural test | Measurement | GFP injected | GluA2 injected | Test statistic | P value |
|---|---|---|---|---|---|
| Elevated plus maze | Speed | 5.7 ± 0.4 cm/s | 5.4 ± 0.4 cm/s | $t_{(21)} = 0.52$ | p = 0.61 |
| | Time in open arms | 11.5 ± 3.7 % | 5.1 ± 1.4 % | $t_{(12.8)} = 1.6$ | p = 0.13 |
| | Open arm entries | 25.3 ± 7.3 | 17.3 ± 3.1 | $t_{(13.6)} = 1.00$ | p = 0.31 |
| Locomotor assay (in a novel photocell activity cage) | Total number of beam breaks | 2908.8 ± 278.9 | 2442.9 ± 158.6 | $t_{(16.0)} = 1.48$ | p = 0.17 |
| | Total number of rearings | 226.8 ± 16.0 | 176.7 ± 13.5 | $t_{(21)} = 2.41$ | p = 0.048* |
| Open field test | Distance travelled | 2299.3 ± 187.8 cm | 2284.0 ± 160.5 cm | $t_{(21)} = 0.07$ | p = 0.94 |
| | Time spent in inner circle | 6.9 ± 1.5 s | 8.1 ± 1.7 s | $t_{(21)} = 0.52$ | p = 0.61 |
| | Latency to enter inner circle | 45.7 ± 9.3 s | 68.6 ± 21.1 s | $t_{(21)} = 0.96$ | p = 0.35 |
| Y-maze spatial novelty preference | Entries into novel arm | 4.45 ± 0.36 | 5.08 ± 0.47 | $t_{(21)} = 1.05$ | p = 0.31 |
| | Novelty preference ratio | 0.60 ± 0.05 | 0.71 ± 0.04 | $t_{(21)} = 1.99$ | p = 0.06 |
| Food neophagia | Latency to first contact | 4.70 [1.50, 9.60] s | 2.55 [2.03, 4.33] s | U = 55.5 | p = 0.53 |
| | Latency to first drink | 13.2 [7.3, 216.4] s | 10.6 [5.0, 48.6] s | U = 54.0 | p = 0.49 |
| Elevated Y-maze spatial reference memory task | Correct choices (Main effect of group) | | | $F_{(1,21)} = 0.01$ | p = 0.92 |
| | Correct choices (Group x session interaction) | | | $F_{(4.1,85.1)} = 0.45$ | p = 0.77 |
| | Success in post-choice baiting | 90.9 ± 4.8 % | 90.8 ± 3.4 % | $t_{(21)} = 0.013$ | p = 0.99 |

**Table S1. Summary of additional behavioural tests.**
The performance on food neophagia is provided as median [inter-quartile range], and tested using Mann-Whitney U test. All other measures represent mean ± SEM, with differences in behaviour assessed using mixed ANOVA or unpaired t tests, with corrections for degrees of freedom depending on sphericity or equality of variance, respectively.
*p<0.05.
